# Long-term in vitro exposure of *Treponema pallidum* to sub-bactericidal doxycycline did not induce resistance: Implications for doxy-PEP and syphilis

**DOI:** 10.1101/2024.06.27.600921

**Authors:** Lauren C. Tantalo, Ann Luetkemeyer, Nicole A. P. Lieberman, B. Ethan Nunley, Carlos Avendaño, Alexander L. Greninger, Connie Celum, Lorenzo Giacani

## Abstract

Doxycycline post-exposure prophylaxis (doxy-PEP) could significantly reduce syphilis incidence. However, the increase in intermittent doxycycline usage might select resistant *Treponema pallidum* (*T. pallidum*) strains. To assess whether resistance to doxycycline could be induced in this pathogen, we exposed the SS14 strain in vitro both intermittently and continuously to a sub-bactericidal doxycycline concentration that still exerts antibiotic pressure. During and after each exposure experiment, we assessed the doxycycline minimal inhibitory concentration in test and control treponemes and performed whole genome sequencing, concluding that no resistance developed. This work suggests that doxycycline-resistant *T. pallidum* is not an immediate threat for doxy-PEP implementation.

## INTRODUCTION

Global syphilis prevalence and incidence are estimated to range between 18-36 million total cases and 5.6-11 million new annual cases, respectively [1, 2]. If untreated, the infection might lead to severe morbidity and, rarely, mortality. Furthermore, vertical transmission of the syphilis agent, *Treponema pallidum subsp. pallidum (T. pallidum*), can result in stillbirth or perinatal death [3]. Syphilis has been resurgent for years in many high-income *nations*. In the U.S., for example, the rate of primary and secondary syphilis reached 17.7 cases of per 100,000 population in 2022, representing a 8.4-fold increase compared to the 2.1 cases per 100,000 population reported in 2000 [4]. A high proportion (29.0%) of the 59,016 primary and secondary syphilis cases reported in 2022 occurred in men who have sex with men (MSM) [4].

A proposed intervention to reverse the increase in syphilis incidence is 200 mg of immediate release doxycycline hyclate or monohydrate within 72 hours after condomless sex to reduce the likelihood of infection with *T. pallidum, Chlamydia trachomatis, and Neisseria gonorrhoeae* [5]. Doxycycline post-exposure prophylaxis (doxy-PEP) reduced the incidence of syphilis, chlamydia, and to a lesser extent gonorrhea in MSM and transgender women [6, 7]. If doxy-PEP is widely adopted, however, the increased and intermittent use of this antibiotic could favor selection of doxycycline (and, in general, tetracyclines) resistance in *T. pallidum*, which has never been confirmed to date, despite reports [13]. Because doxycycline is an effective second-line antibiotic, particularly useful for penicillin-allergic individuals, the spread of resistant *T. pallidum* strains would have substantial consequences in the management of syphilis patients.

We conducted in vitro studies to assess whether genetic resistance to doxycycline could be induced in *T. pallidum* by a) determining a sub-treponemicidal concentration of doxycycline that could exert detectable antibiotic pressure on the pathogen in vitro, b) exposing *T. pallidum* to doxycycline intermittently over seven months, and continuously for 10 weeks, and c) assessing changes in the *T. pallidum* genome and in the doxycycline minimal inhibitory concentration (MIC) for the pathogen after each intermittent exposure or at the end of the continual exposure experiment.

## METHODS

### *T. pallidum* strain and propagation

The *T. pallidum* SS14 strain, isolated in 1977, was kindly provided by Dr. Sandra A. Larsen (CDC, Atlanta, GA). Strain propagation was performed per established protocols [9]. For the exposure experiments, doxycycline (Millipore-Sigma) was added to the culture media from a 5.0 µg/ml stock solution in sterile water to the desired final concentration.

The initial determination of the doxycycline MIC for *T. pallidum* was performed as reported [10] by incubating *T. pallidum* in vitro for seven days in concentrations ranging from 10 ng/ml to 600 ng/ml in 8 replicate wells of a 96-well plate (each containing 5×10^4^ *T. pallidum* cells and 3×10^3^ Sf1Ep rabbit cells), followed by determination of treponemal burden by qPCR in test and no-antibiotic wells.

To further define a sub-bactericidal concentration of doxycycline able to exert antibiotic pressure for the long-term exposure experiments, we performed a “limited exposure “ experiment, where *T. pallidum* was grown in vitro in 100 ng/ml of doxycycline for one or two weeks, respectively, followed by transfer to antibiotic-free media to assess whether cultures would recover.

For the “intermittent exposure “ experiment, *T. pallidum* was treated with 50 ng/ml doxycycline (selected based on the results of the experiments mentioned above) for two consecutive weeks followed by a two- to three-week recovery period, and for a total of six exposures over a seven-month period of continuous cultivation. This protocol allowed the culture to rebound for recovery of enough treponemes to assess changes in susceptibility to doxycycline and perform WGS after each exposure. For the ten-week “continuous exposure “ experiment, *T. pallidum* was treated with doxycycline at 10, 20, 30, 40, 50, and 100 ng/ml. Unexposed control treponemes were propagated in parallel throughout both experiments. Due to the necessity of counting treponemes, the continuous and intermittent exposure experiments were conducted in 6-well culture plates, with media, treponemal and Sf1Ep inocula adjusted based on the well area.

Each week, treponemes were recovered from culture plates and enumerated by darkfield microscopy (DFM). A total of 10^7^ T. pallidum cells were then transferred into the wells of a newly prepared plate. The sub-culturing/enumeration process was repeated the following week. If, due to antibiotic pressure, a week ‘s culture exhibited low or no quantifiable (by DFM) treponemal yield such that fewer (or an unknown number of) organisms could be re-inoculated, then 500 µl of culture extract (i.e., the maximum volume transferable into a single well of a new 6-well plate) were inoculated instead. At each weekly passage, culture samples were also collected by centrifugation at 20,000 RCF for 10 min, and pelleted cells were resuspended in either 200 µl of lysis buffer (10mM Tris, 0.1M EDTA, and 1% SDS) or 400 µl of Trizol (Thermo Fisher) for DNA or RNA extraction, respectively.

### Nucleic acid extraction, amplification, and whole genome sequencing

DNA was extracted either as previously described [10] for experiments conducted using 96-well plates (for MIC determination), or using the QIAmp DNA Mini Kit (Qiagen) according to the manufacturer ‘s instructions for individual samples for whole genome sequencing (WGS). RNA was isolated according to the Trizol manufacturer ‘s instructions and treated with the TURBO DNA-free kit (Thermo Fisher) to eliminate residual DNA. DNA-free RNA was reverse transcribed using the SuperScript III kit (Thermo Fisher). Qualitative amplification of template cDNA targeting the *tp0574* gene were performed as reported [10].

Library preparation and enrichment of *T. pallidum* DNA ahead of WGS, and the pipeline for genomic data analysis were previously published [11] and are briefly reported in the Supplementary Data File. Consensus sequences and FATQ data are available in BioProject PRJNA1113685.

## RESULTS

In vitro exposure of *T. pallidum* to doxycycline supported that the MIC for the syphilis agent, defined as the lowest antibiotic dilution at which the tp0574 qPCR value was significantly lower than the positive control (treponemes grown for 7 days in absence of antibiotic), was 50 ng/ml, although this concentration only partially inhibited growth over a 7-day exposure (Fig.S1A). Based on our definition of minimal bactericidal concentration (MBC) as the lowest concentration at which there is no bacterial growth after sub-culturing of exposed bacteria into antibiotic-free media, 300 ng/ml of doxycycline were necessary (Fig.S1B). The “limited exposure “ experiment confirmed that exposure of *T. pallidum* to 100 ng/ml for one or two weeks was not completely bactericidal in vitro, as all cultures recovered over time (Fig.S2). Based on these results, a concentration of 50 ng/ml was selected for the “intermittent exposure “ experiment, which would result in measurable antibiotic pressure, and ensure culture recovery following a 2-week exposure.

Compared to control treponemes not exposed to doxycycline, treponemal numbers dropped during each exposure, to always recover after withdrawing treatment (Fig.1A). During the last two exposure cycles however, overall cell numbers decreased less markedly, and recovered faster to the level of control treponemes (Fig.1A). Doxycycline MIC determination for control and treated cultures after exposures 1-5 (Fig.S3), as well as after the sixth and final exposure (Fig.1B-D), however, did not support an increase in the doxycycline MIC for *T. pallidum*. Whole-genome sequencing of the test and control strain after each exposure did not yield evidence of fixation of mutations related to resistance to doxycycline in the treated strain based on analysis by the Comprehensive Antibiotic Resistance Database (CARD) software (https://card.mcmaster.ca/) (TableS1/S2). No treatment-emergent mutations in 16S rRNA were recovered at 1% allele frequency across all doxycycline-selected isolates. Progressive fixation of a non-synonymous SNP (resulting in the T46I amino acid change) in TP0319-TmpC (Fig.S4), a substrate-binding periplasmic lipoprotein, was seen during both the intermittent and continuous exposure experiments. Based on its presence at low level in the inoculum, as well as its enrichment also during longitudinal culture of untreated controls, we concluded that this was not a mutation conveying resistance but related to culture adaptation of the SS14 strain.

**Figure 1.**
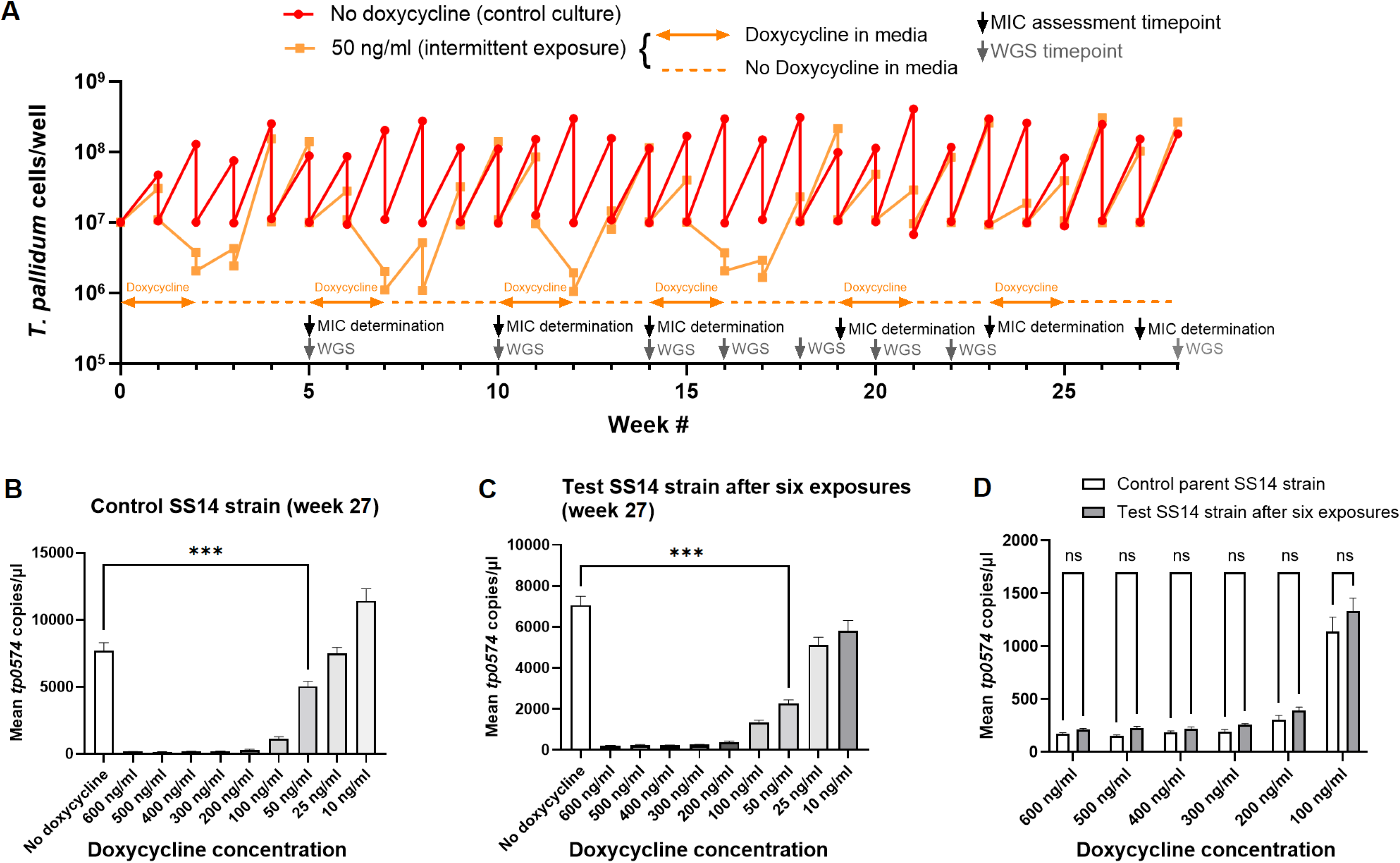
**(A)** *T. pallidum* growth in vitro when exposed to pulses of 50 ng/ml doxycycline for 2 weeks, as indicated by orange bars 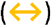, followed by a 2/3 week-long recovery period in absence of antibiotic 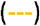 for a total of 28 weeks. Concentrations were determined by DFM. Arrows indicate dates of MIC (↓) and WGS 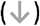 assays performed on recovered cultures. MIC assays on control treponemes (B) and test treponemes (C) harvested at week 27, after six pulsed exposures to the test strain to doxycycline. (D) Comparison of treponemal growth (or lack thereof) in different doxycycline concentration of control (white bars) and test treponemes (grey bars) following sux pulsed exposures to doxycycline. Analyses were performed using ANOVA or T-test with significance set at p ≤0.05.

The results of the “continuous exposure “ experiment showed that 40, 50, and 100 ng/ml affected *T. pallidum* growth in a dose dependent manner, while 10, 20 and 30 ng/ml did not appear to induce measurable (by DFM) pressure on the pathogen based on culture yields (Fig.2A). Amplification of *T. pallidum* tp0574 RNA as a surrogate of viability (Fig.2B), demonstrated that approximately 4 weeks of continuous exposure to 50 ng/ml of doxycycline were necessary for loss of amplification signal, while between seven to 14 days were necessary when *T. pallidum* was exposed to 100 ng/ml (Fig.2B). *T. pallidum* cells exposed to 40 ng/ml of doxycycline were not quantifiable by DFM after four weeks (Fig.2B), but the message amplification signal was detectable at every subculturing event during the 10-week duration of the experiment (Fig.2B). Compared to RNA, DNA signal (Fig.2C) persisted longer. At the end of the experiment, the sequenced genomes of treponemes propagated in 10, 20, or 30 ng/ml did not present evidence of mutations related to doxycycline resistance (TableS1/S2) and repeating the MIC assessment was deemed unnecessary.

**Figure 2.**
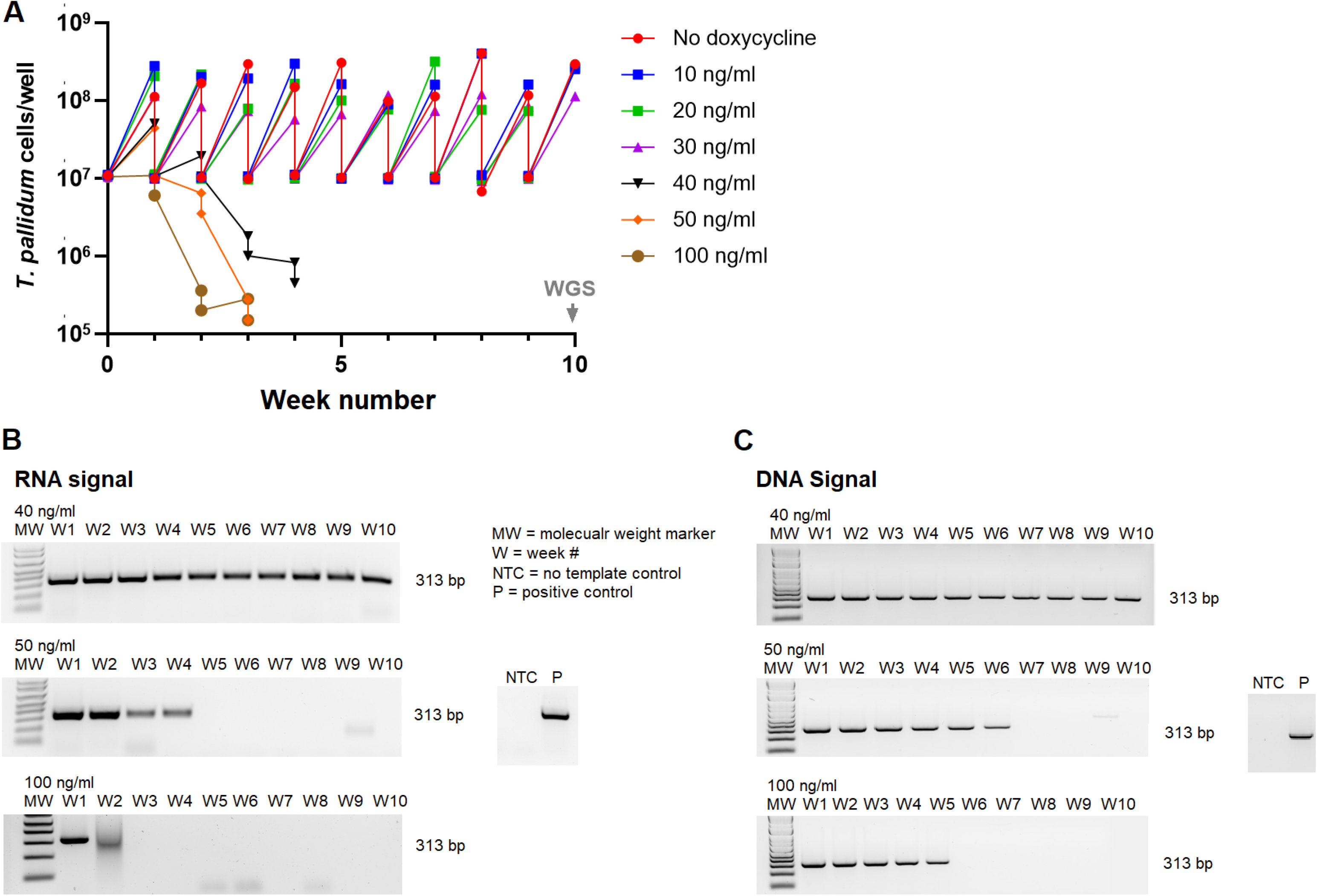
**(A)** *T. pallidum* growth in vitro when continually exposed to concentrations of doxycycline between 10 and 100 ng/ml for 7 days, or in absence of antibiotic. The arrow 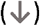 indicates the date of WGS assays performed on recovered cultures at the end of the 10-week experiment. (B) Qualitative Reverse Transcription-PCR of *T. pallidum* samples harvested at every subculturing event of treponemes propagated in presence of 40 ng/ml of doxycycline (top photograph), 50 ng/ml (middle photograph), and 100 ng/ml (bottom photograph) following RNA extraction and cDNA synthesis. (C) Qualitative amplification of *T. pallidum* samples harvested at every subculturing event of treponemes propagated in presence of 40 ng/ml of doxycycline (top photograph), 50 ng/ml (middle photograph), and 100 ng/ml (bottom photograph) following DNA extraction showing longer persistence of DNA signal in culture compared to RNA signal. MW: molecular weight marker; W: week # (harvest treponemes); NTC: no template control; P: positive control (*T. pallidum* DNA). The amplification target (*tp0574)* is visible as a 313 bp band.

## DISCUSSION

The use of macrolides as an oral alternative to parenteral penicillin resulted almost all circulating *T. pallidum* strains being resistant to macrolides in the U.S. [8]. Because the use of doxycycline for syphilis prevention is expected to increase significantly based on high efficacy of doxy-PEP in several clinical trials among MSM [6, 7], the possibility that *T. pallidum* might develop resistance to this antibiotic needed to be assessed. Macrolide resistance in *T. pallidum* is due to either the A2058G or A2059G point mutation in the 23S rRNA gene of the pathogen [8]. Both macrolides and tetracyclines are protein synthesis inhibitors, and although they act on different targets and subunits of the bacterial ribosome, resistance to doxycycline could also result from single point mutations in the *T. pallidum* 16S rRNA genes or specific ribosomal proteins [12]. Our in vitro experiments did not show evidence of such changes in treponemes exposed to sub-bactericidal doxycycline; susceptibility to doxycycline did not decrease compared to the unexposed parent strain.

*T. pallidum* has limited genome plasticity and no plasmids [11], therefore the likelihood that resistance genes encoding efflux pumps, ribosomal protection proteins, or proteins for enzymatic inactivation could be stably acquired from environmental sources is minimal. A recent report, however, mentioned the discovery of transitions in a mutational hotspot for doxycycline resistance, namely the TGA triplet at position 965-967 of the 16S rRNA gene, in a subset (4/544) of available *T. pallidum* genomes [13]. At this time, however, whether these changes would confer resistance in *T. pallidum* is unclear, given that no isolate carrying these mutations is available for testing and no information is provided concerning whether these genomes were recovered from patients that failed doxycycline treatment. These data highlight the need for isolation of clinical *T. pallidum* strains to perform phenotypic testing and for using genetic engineering of *T. pallidum* to characterize potential resistance mutations when isolates do not exist [14].

A limitation of this study is that a single *T. pallidum* strain was used due to the significant workload associated with the cultivation procedures. Even though *T. pallidum* is genetically monomorphic [12], repeating the experiments with additional more recently isolated strains could be informative. Another priority is to perform longer intermittent and continuous exposure experiments with lower doxycycline concentrations than used here as might be observed with less frequent dosing of doxy-PEP.

Haaland et al. [15] reported that following a single 200 mg doxycycline dose, mean plasma concentrations did not fall below 100 ng/ml for 2.9 and 3.6 days, respectively, in plasma and rectal secretions. Although we did not conduct exposure experiments using >100 ng/ml of doxycycline, one could postulate that T. pallidum killing by concentrations >100 ng/ml would be rapid, explaining doxy-PEP efficacy.

In summary, we did not derive a doxycycline-resistant strain with subtherapeutic doses of doxycycline in vitro, nor did we recover mutations in putative doxycycline resistance related genes in doxycycline-exposed isolates. Although our findings do not preclude that resistance could eventually develop in clinical settings depending on the prevalence and persistence of doxy-PEP use, our data suggest that the development of resistance to doxycycline in T. pallidum is not an immediate concern.

## FOOTNOTES

This work was supported by NIAID grant R01AI143439-05 to A.L. and C.C.

All authors declare no conflict of interest

These data have been in part presented at the NIAID workshop “*Accelerating development of alternative therapies to benzathine penicillin for the treatment of syphilis “*. February 13-14, 2024

The authors are grateful to Dr. Richard Haaland, Research Microbiologist at Centers for Disease Control and Prevention (Atlanta, GA) for critical review of the manuscript data.

## Notes

### Competing Interest Statement

The authors have declared no competing interest.

## REFERENCES

1. WHO, Prevalence and incidence of selected sexually transmitted infections Chlamydia trachomatis, Neisseria gonorrhoeae, syphilis and Trichomonas vaginalis: methods and results used by WHO to generate 2005 estimates. World Health Organization, Geneva, 2011.

2. Gerbase, A.C., J.T. Rowley, and T.E. Mertens, Global epidemiology of sexually transmitted diseases. The Lancet, 1998. 351.

3. LaFond, R.E. and S.A. Lukehart, Biological basis for syphilis. Clin Microbiol Rev, 2006. 19(1): p. 29–49.

4. CDC, 2022 Sexually Transmitted Disease Surveillance. Atlanta, GA: U.S. Department of Health and Human Services: Centers for Disease Control and Prevention, 2024.

5. Lh, B., et al., CDC Clinical Guidelines on the Use of Doxycycline Postexposure Prophylaxis for Bacterial Sexually Transmitted Infection Prevention, United States, 2024. MMWR Recomm Rep, 2024. 73(No. RR-2):1–8.

6. Luetkemeyer, A.F., et al., Postexposure Doxycycline to Prevent Bacterial Sexually Transmitted Infections. N Engl J Med, 2023. 388(14): p. 1296–1306.

7. Molina, J.M., et al., Post-exposure prophylaxis with doxycycline to prevent sexually transmitted infections in men who have sex with men: an open-label randomised substudy of the ANRS IPERGAY trial. Lancet Infect Dis, 2018. 18(3): p. 308–317.

8. Stamm, L.V., Global challenge of antibiotic-resistant Treponema pallidum. Antimicrob Agents Chemother, 2009. 54(2): p. 583–9.

9. Edmondson, D.G., B. Hu, and S.J. Norris, Long-Term In Vitro Culture of the Syphilis Spirochete Treponema pallidum subsp. pallidum. mBio, 2018. 9(3).

10. Tantalo, L.C., et al., Antimicrobial susceptibility of Treponema pallidum subspecies pallidum: an in-vitro study. Lancet Microbe, 2023. 4(12): p. e994–e1004.

11. Lieberman, N.A.P., et al., Treponema pallidum genome sequencing from six continents reveals variability in vaccine candidate genes and dominance of Nichols clade strains in Madagascar. PLoS Negl Trop Dis, 2021. 15(12): p. e0010063.

12. Nguyen, F., et al., Tetracycline antibiotics and resistance mechanisms. Biol Chem, 2014. 395(5): p. 559–75.

13. Manoharan-Basil, S., et al., Tetracycline Resistance Genes in Treponema spp.-an Analysis of 4355 Spirochaetales Genomes, in Preprints. 2024, Preprints.

14. Romeis, E., et al., Genetic engineering of Treponema pallidum subsp. pallidum, the Syphilis Spirochete. PLoS Pathog, 2021. 17(7): p. e1009612.

15. Haaland, R.E., et al., Pharmacokinetics of single dose doxycycline in the rectum, vagina, and urethra: implications for prevention of bacterial sexually transmitted infections. EBioMedicine, 2024. 101: p. 105037.

