## Supplemental material for "Long-term in vitro exposure of *Treponema pallidum* to sub-bactericidal doxycycline did not induce resistance: Implications for doxy-PEP and syphilis"

### ***In vitro* cultivation of *T. pallidum***

Procedures for the *in vitro* cultivation of *T. pallidum* were conducted as described previously [1] Briefly, two sets of cultures were prepared: one for the susceptibility assay and one for the bactericidal/recovery assay. The susceptibility assay involved testing drug concentration in 96-well plates (Corning Inc., 8x12 format). Each drug concentration was treated as a separate experimental group and tested eight times in eight replicate wells. A control group without antibiotics was included, also tested in eight replicate wells.

Wells were seeded with 3x10^3^ rabbit Sf1Ep cells in 150 μl MEM media and incubated overnight. The next day, 150 μl TpCM2 media (equilibrated overnight in a 34°C tri-gas incubator) was added for three hours to acclimate cells to low oxygen. After removing TpCM2, 148.5 μl of a 3.3x10^5^ *T. pallidum* cells/ml inoculum was added to each well (5x10^4^ *T. pallidum*/well). Antibiotic solutions (1.5 µl) were added from 100X concentrated stocks without altering volume followed by incubation at 34ºC in a tri-gas incubator until harvest. Experimental and control wells with varying antibiotic concentrations were harvested after a week for DNA quantification. Treponemal burden was assessed using qPCR targeting the *T. pallidum*-specific *tp0574* gene (sensitivity of ~10 treponemal genomes per reaction).

A second set of plates was prepared for the bactericidal/recovery assay. Treponemes exposed to the drug concentration were sub-cultured into antibiotic-free recovery plates. These plates were incubated for seven more days before DNA extraction.

**Outcomes and statistical analyses**

The MIC for each experiment was defined as the lowest antibiotic dilution at which the *tp0574* qPCR values were significantly lower than the positive control (Day-7 control group), which is consistent with the broth dilution procedure and was used previously by Haynes *et al* [2]. The minimum bactericidal concentration (MBC) was defined as the lowest concentration at which there was no bacterial growth after sub-culturing into the antibiotic-free media.

The sample size consisted of 8 replicates per drug concentration and control group. Each sample represented a technical replicate from the same source mixture. Rigorous experimental conditions minimized significant variations among samples.

For statistical analyses, we employed Analyses were performed using ANOVA or t-test with significance set at p≤0.05 to compare the mean qPCR values among groups with different antibiotic concentrations and control group. Dunn's test was used for pairwise comparisons between specific groups and control groups at Day 7.

**Whole genome sequencing**

For WGS, pre-capture libraries were prepared from up to 100 ng input DNA using the KAPA Hyperplus kit (Roche) and TruSeq adapters and barcoded primers (Illumina), following the manufacturer’s protocols, yielding an average fragment size >500 bp. Hybrid capture of *T. pallidum* DNA was performed overnight (>16 hours) using a custom IDT xGen panel designed against the reference NC_ 010741 genome, following the manufacturer’s protocol. Libraries were sequenced on a Nextseq 2000 (Illumina). Paired end 2x151 bp reads were trimmed using Trimmomatic v0.39 and mapped using Bowtie2 to the SS14 reference (NC_021508.1). Manual confirmation of expected coverage and junctions was performed by visual inspection in Geneious Prime v2020.1.2. The detailed analytical pipeline for genome sequencing was previously reported [3]. Consensus sequences are available in the NCBI SRA in BioProject PRJNA1113685 “In vitro doxycycline treatment of *T. pallidum*”.

**Figure S1**

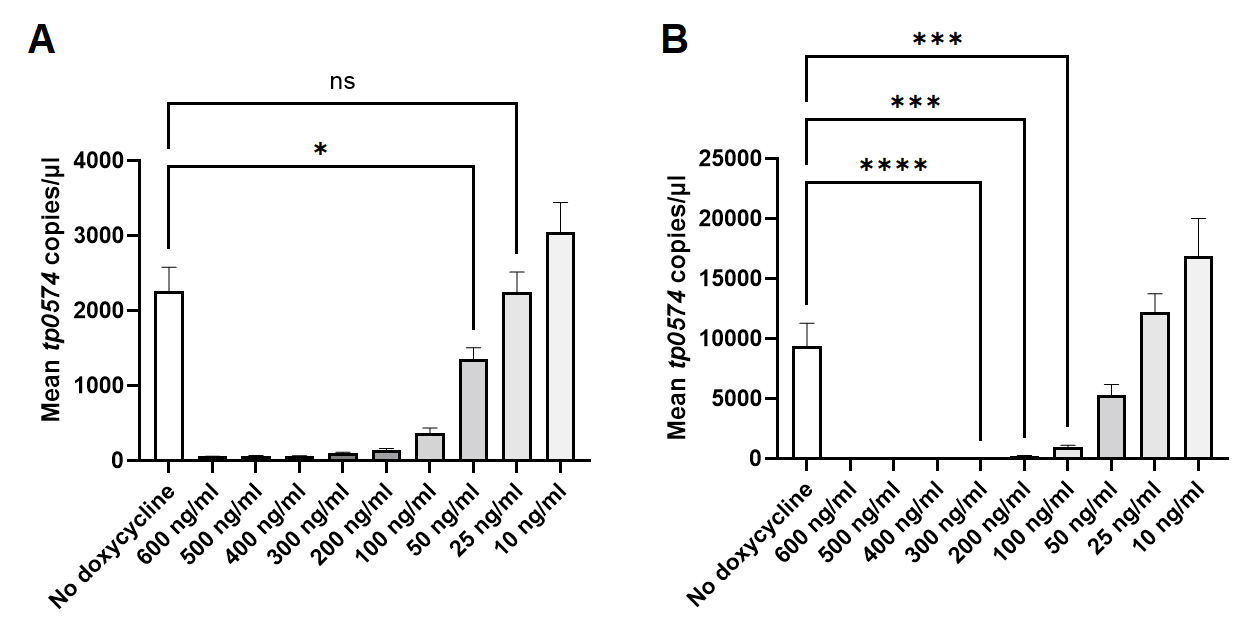

**Figure S1**. **(A)** *T. pallidum* growth *in vitro* when exposed to concentrations of doxycycline between 10 and 100 ng/ml for 7 days, and in absence of antibiotic. The MIC (50 ng/ml) was defined as the lowest antibiotic concentration at which growth (measured by DNA quantification) was significantly lower than the positive control (No doxycycline well). **(B)** Recovery assays in no-antibiotic plate of *T. pallidum* exposed to 10-600 ng/ml of doxycycline shown in Fig.1A to determine the doxycycline MBC, defined as the lowest concentration at which there is no bacterial growth after sub-culturing of exposed bacteria into antibiotic-free media. Values represent mean *tp0574* gene copies per unit of volume. Analyses were performed using ANOVA or t-test with significance set at p≤0.05.

**Figure S2**

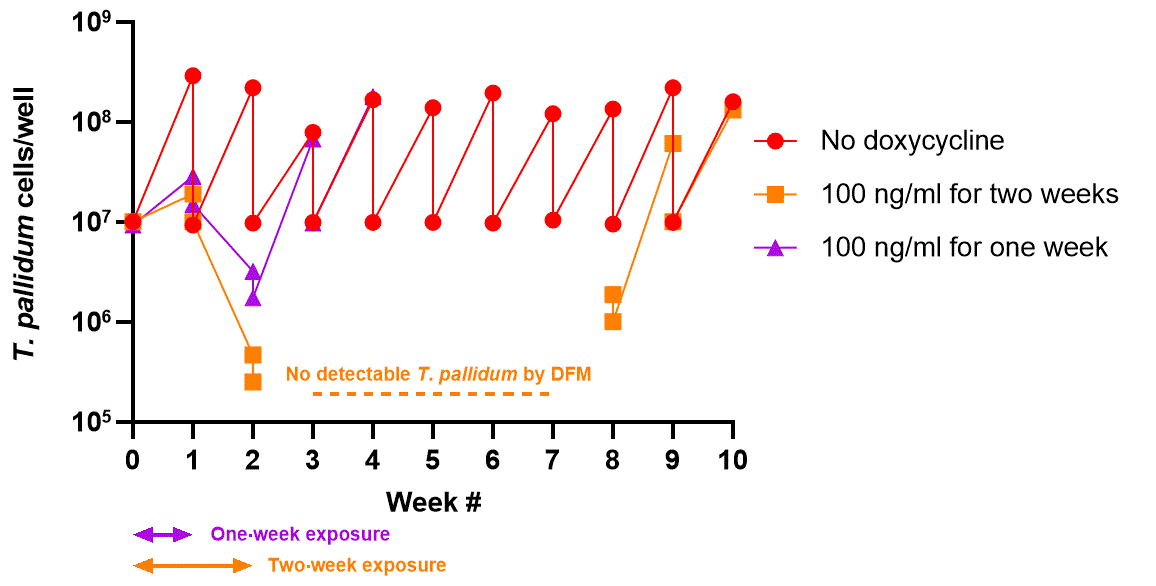

**Figure S2.** Exposure of *T. pallidum* cultures to 100 ng/ml of doxycycline for one (**↔**) or two (**↔**) consecutive weeks (“limited exposure” experiment). *In vitro* growth of a *T. pallidum* inoculum exposed to 100 ng/ml of doxycycline for one week (purple triangles) is inhibited compared to the control unexposed culture (red dots), but the culture recovers to the level of the control culture in approximately three weeks (Week 4) after doxycycline withdrawal. The growth of the same *T. pallidum* inoculum exposed to 100 ng/ml of doxycycline for two weeks (orange squares) is more severely inhibited, and cells become uncountable by DFM for five weeks (Week 3-Week7; **----**) after doxycycline withdrawal. Following eight sub-culturing events (five of which were blind to the number of inoculated treponemes), the exposed culture recovered to the level of control (untreated) treponemes by week #10.

**Figure S3**

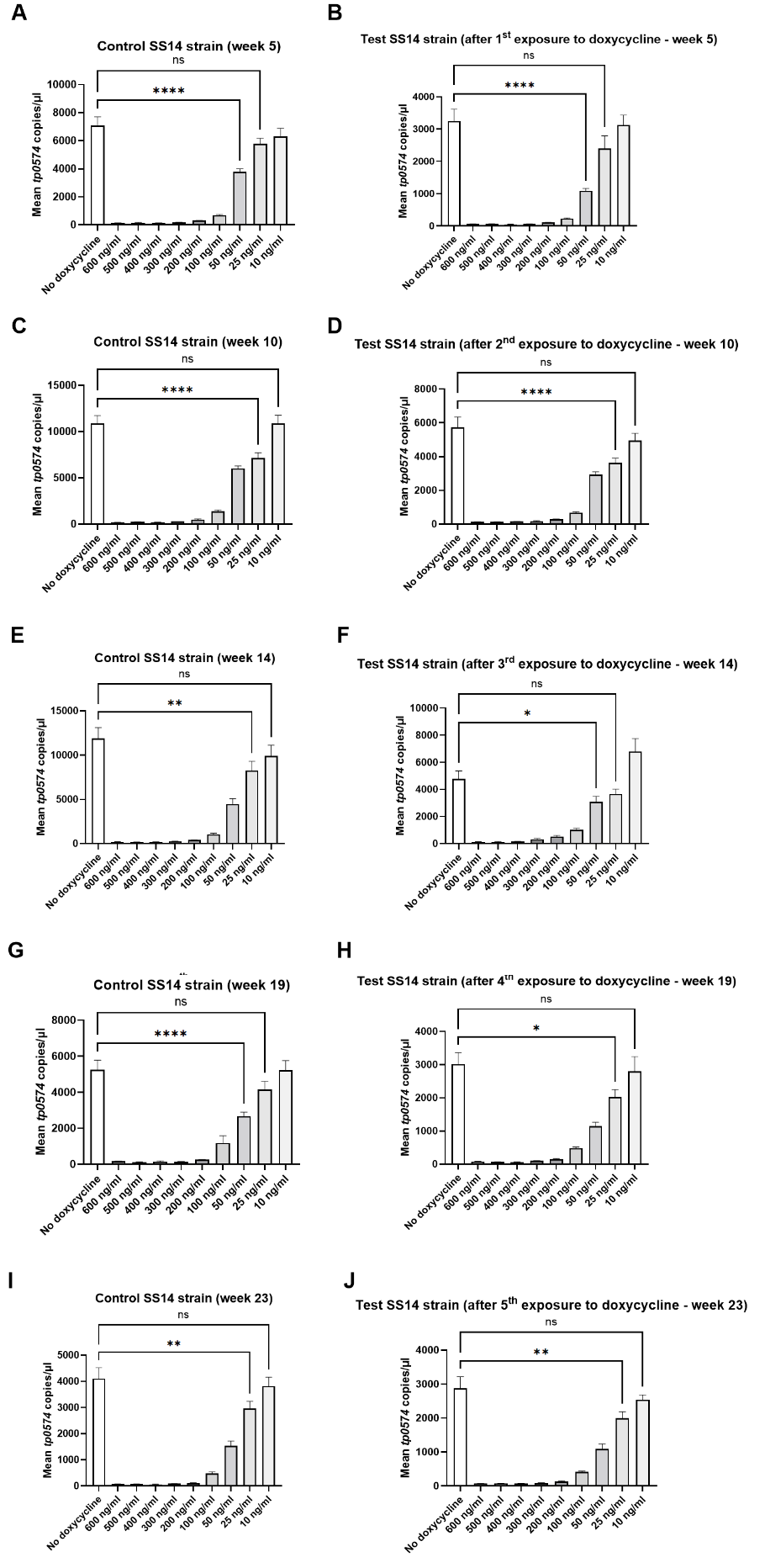

**Figure S3. (A-J)** Growth in presence of doxycycline (10-100 ng/ml for 7 days) of *T. pallidum* control strain and test strain after each exposure to doxycycline (50 ng/ml for two weeks), supporting no change in the pathogen MIC for doxycycline as the result or repeated exposures.

**Figure S4**

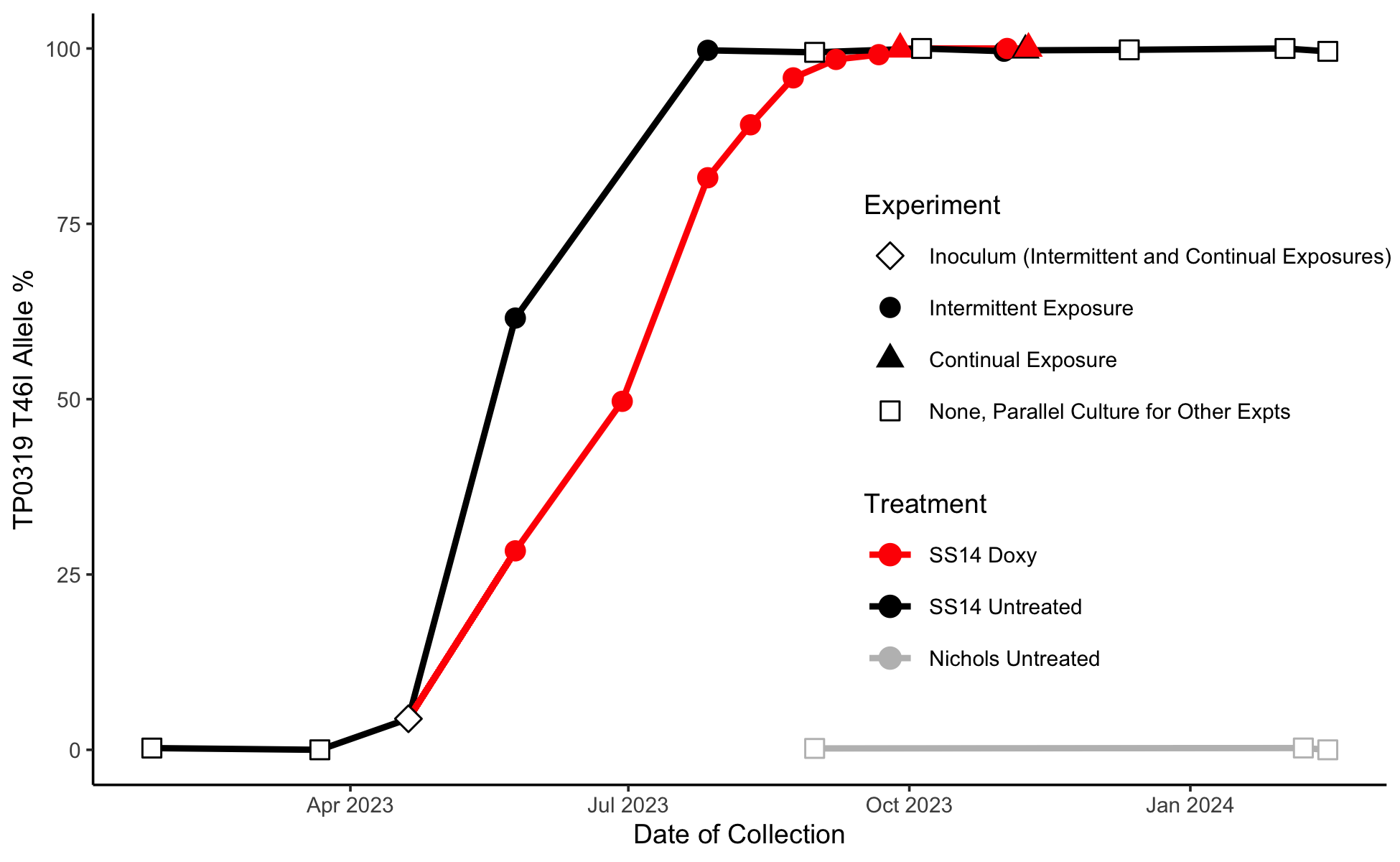

**Figure S4.** Fixation of the T46I mutation in SS14 strain propagated *in vitro* in presence of sub-bactericidal doxycycline (red) and in unexposed control strains propagated in parallel during the intermittent and continuous exposure experiment, respectively. Fixation of T46I did not occur in the Nichols strain propagated in parallel to the SS14 for other experimental purposes.

**Table S1.** Analysis of all sequenced genomes from test strains following intermittent or continuous exposure to doxycycline (Fig.1,2), or in control strains. Allele frequencies per sample for all SNPs that had at least one sample at > 10% minor allele frequency. Columns are named with info including nt position on NC_02150801, locus tag, gene name, and protein effect.

**Table S2.** Results of the analysis of all sequenced genomes using the Comprehensive Antibiotic Resistance Database (CARD; https://card.mcmaster.ca/) supporting that no genes conferring resistance to doxycycline could be identified in test strain genomes following intermittent or continuous exposure to doxycycline (Fig.1,2), or in control strains.

**References**

1.	Edmondson DG, Hu B, Norris SJ. Long-Term In Vitro Culture of the Syphilis Spirochete *Treponema pallidum* subsp. *pallidum*. mBio. 2018;9(3). Epub 2018/06/28. doi: 10.1128/mBio.01153-18. PubMed PMID: 29946052; PubMed Central PMCID: PMCPMC6020297.

2.	Haynes AM, Giacani L, Mayans MV, Ubals M, Nieto C, Pérez-Mañá C, et al. Efficacy of linezolid on *Treponema pallidum*, the syphilis agent: A preclinical study. EBioMedicine. 2021;65:103281. Epub 2021/03/16. doi: 10.1016/j.ebiom.2021.103281. PubMed PMID: 33721817; PubMed Central PMCID: PMCPMC7973135.

3.	Lieberman NAP, Lin MJ, Xie H, Shrestha L, Nguyen T, Huang ML, et al. *Treponema pallidum* genome sequencing from six continents reveals variability in vaccine candidate genes and dominance of Nichols clade strains in Madagascar. PLoS Negl Trop Dis. 2021;15(12):e0010063. Epub 2021/12/23. doi: 10.1371/journal.pntd.0010063. PubMed PMID: 34936652.
